# Proximity proteomics of primary cilia in human hypothalamic neurons

**DOI:** 10.1101/2025.05.11.653368

**Authors:** Viviana Macarelli, Tommy J. Sroka, Jan N. Hansen, Ulrika Axelsson, David U. Mick, Florian T. Merkle

## Abstract

Primary cilia are hair-like sensory organelles that project from the cell bodies of most cell types, including appetite-regulatory hypothalamic neurons where they likely help sense metabolic factors to regulate food intake. We hypothesized that characterising the proteins present in the primary cilia of hypothalamic neurons would shed mechanistic insights into their sensory role and identify new therapeutic targets for obesity. We therefore targeted the ascorbate peroxidase APEX2 to primary cilia in human induced pluripotent stem cells (hiPSC)-derived hypothalamic neurons to biotinylate and identify ciliary proteins. Among the cilia-enriched proteins, we identified synaptic proteins, neurotransmitter receptors, and cell-cell adhesion and axon guidance proteins, extending recent findings that primary cilia interact with neuronal synapses. We also found genes associated with increased body weight and metabolic phenotypes that could represent new therapeutic targets including the lysophosphatidic receptor 1 (LPAR1), which we validated is cilia-localized and we confirmed that its ligand (LPA) mediates ciliary shortening. These findings provide insights into the molecular mechanisms by which primary cilia functionally impact appetite-regulatory neurons.

## Introduction

The primary cilium is a sensory organelle protruding from the surfaces of most mammalian cells that is thought to serve as a cellular antenna (Singla and Reiter 2006). The ciliary membrane is enriched with certain G protein-coupled receptors (GPCRs) that sense extracellular cues and transmit them to the cell body, making cilia essential for normal development and tissue homeostasis (Derderian, Canales, and Reiter 2023). Indeed, ciliary dysfunction can lead to syndromic disorders, known as ciliopathies, which affect different organs and cell-types (Mill, Christensen, and Pedersen 2023).

In brain development, primary cilia regulate neuronal progenitor cell (NPC) proliferation, differentiation and migration, but they are also found in sensory and central nervous system (CNS) neurons of the adult brain (Jurisch-Yaksi, Wachten, and Gopalakrishnan 2024; Suciu and Caspary 2021). Their dysfunction in brain cells is linked to neurodevelopmental (S. Liu et al. 2021), neurodegenerative (Ma et al. 2022) and psychiatric disorders (Ma et al. 2022; Alhassen et al. 2021). Primary cilia of adult CNS neurons can sense neurotransmitters and other signals and may regulate synaptic function and connectivity (Jurisch-Yaksi, Wachten, and Gopalakrishnan 2024). Indeed, primary cilia can form synapses with serotonergic neurons (Jurisch-Yaksi, Wachten, and Gopalakrishnan 2024; Sheu et al. 2022) and are thought to help regulate the excitability of interneurons, hippocampal neurons, pyramidal neurons and cortical neurons (J. Guo et al. 2017; Jang et al. 2023; Tereshko et al. 2021; Vien et al. 2023; Yang et al. 2025). However, the molecular mechanisms by which primary cilia regulate neuronal function in the adult brain are still poorly understood.

One brain region of particular interest is the hypothalamus, where primary cilia are likely to be important in the regulation of appetite, energy expenditure, and body weight (Brewer et al. 2022). Indeed, the ciliopathies Bardet-Biedl syndrome (BBS), Alström syndrome (ALMS), Carpenter syndrome, and MORM syndrome, alongside other features (Brewer et al. 2022), are associated with obesity; a highly heritable and largely-brain-based condition (Brewer et al. 2022; NCD Risk Factor Collaboration 2017; Locke et al. 2015) often defined by a body-mass-index (BMI) of over 30 kg/m2 (Brewer et al. 2022; NCD Risk Factor Collaboration 2017). Targeted deletion of genes required for normal ciliary function, such as intraflagellar transport 88 (*Ift88*), kinesin family member 3A (*Kif3a*), or Bardet-Biedl syndrome 1 (*Bbs1*), in hypothalamic neurons is sufficient to drive increased food intake and body weight in mice (Davenport et al. 2007; D.-F. Guo et al. 2019).

Similarly, receptors with known roles in feeding behavior, such as the melanocortin-4 receptor (Mc4r), melanin-concentrating hormone receptor 1 (Mchr1), the somatostatin receptor type 3 (Sstr3), the neuropeptide Y receptor 2 (Npy2r), and the G protein–coupled receptor 75 (Gpr75) normally localize to hypothalamic cilia but many of them fail to do so in BBS mouse models (Berbari et al. 2008; Loktev and Jackson 2013; Jiang, Xun, and Zhang 2024). Ablation of cilia in Mc4r+ neurons of the paraventricular nucleus of the hypothalamus (PVN) causes obesity (Wang et al. 2021), and the ciliary localisation of Mc4r is dynamically regulated by feeding and fasting (Ojeda-Naharros et al. 2025). However, other mechanisms by which primary cilia might sense metabolic signals to regulate energy homeostasis in the PVN and other hypothalamic nuclei remain poorly understood, largely due to a lack of scalable discovery tools to characterise the ciliary proteome in target cell populations.

The sensory function of primary cilia is closely linked to the receptors and proteins that are present within these organelles, which influence the signals sensed and the downstream response. To the best of our knowledge the only reported characterization of the protein content of neuronal cilia has been done in the mouse forebrain (Loukil et al. 2023) and in the dorsal and ventral regions of the mouse embryonic brain (X. Liu et al. 2025). Here, we used APEX2-mediated proximity proteomics (Shkel et al. 2022) of cultures of predominantly appetite-regulatory neurons of the hypothalamic arcuate nucleus (ARC). We identified 141 significantly enriched proteins in hypothalamic primary cilia including well-known ciliary proteins and many others whose ciliary association is novel. These neuronal ciliary proteins are thought to mediate cell-cell contact, axon guidance, synapse function, and neurotransmitter and neuropeptide sensing. Notably, we confirmed the ciliary localization of the obesity-associated protein lysophosphatidic acid receptor 1 (LPAR1) and demonstrated that its ligand LPA significantly shortened the length of primary cilia in hypothalamic cells. These findings reveal insights into the function of human neuronal primary cilia, and identify a candidate obesity-associated ciliary gene that may mechanistically contribute to appetite regulation and obesity.

## Methods

### Reagents and resources

| REAGENT or RESOURCE | SOURCE or DESCRIPTION | CATALOG |
| --- | --- | --- |
| **Chemicals, peptides, and recombinant proteins** |  |  |
| 100x Protease inhibitor cocktail | Sigma-Aldrich | P8849 |
| 10x Tris/Glycine/SDS running buffer | BioRad | 161-0732 |
| 16% Formaldehyde | ThermoFisher Scientific | 28908 |
| 4-12% Bis-Tris gel | Invitrogen | NP0321 |
| Ascorbic acid | Sigma Aldrich | A4403 |
| B27 supplement | ThermoFisher Scientific | 17504001 |
| Biotin-phenol | Iris Biotech | LS-3500.0250 |
| Calcium chloride solution | Merck | 21115 |
| CHIR99021 | Cambridge Bioscience | SM13 |
| Chloroacetamide | Sigma-Aldrich | C0267 |
| DAPT | DNSK International | DNSK-EI-01 |
| D-Biotin | ThermoFisher Scientific | B20656 |
| Dimethyl sulfoxide (DMSO) | Sigma-Aldrich | D2650 |
| DMEM/F12 with GlutaMAX | ThermoFisher Scientific | 31331093 |
| Ethylenediaminetetraacetic acid (EDTA) | Sigma-Aldrich | 3690 |
| GABA | Tocris | 344 |
| Geltrex Matrix | ThermoFisher Scientific | A1413202 |
| Glutamax | ThermoFisher Scientific | 35050038 |
| Hydroxylamine solution 5% | Sigma | 438227 |
| Human BDNF | Qkine | Qk050 |
| Human GDNF | PeptoTech | 450-10 |
| Human NT-3 | Cambridge Bioscience | GFH99 |
| Laminin | Sigma Aldrich | L2020 |
| Laminin 511 | Takara bio | T304 |
| OASIS HLB 96-well μElution Plate | Waters | 186001828BA |
| Oleoyl-L-α-lysophosphatidic acid sodium salt | Sigma | L7260 |
| LICOR destaining solution | LICOR | 928-11090 |
| LICOR Revert Total Protein Stain | LICOR | 926-11011 |
| LICOR TBS blocking solution | LICOR | 927-50000 |
| LICOR wash solution | LICOR | 928-40002 |
| LM22A4 | Tocris | 4607/5 |
| MEM Non-Essential amino acids | ThermoFisher Scientific | 11140035 |
| Mobicols mini spin columns | MoBiTec | M105135F |
| N-2 Supplement | ThermoFisher Scientific | 17502048 |
| Neurobasal- A | ThermoFisher Scientific | 10888022 |
| NKH 447 | Sigma Aldrich | N3290 |
| Normal Donkey serum | Strattech Scientific | 017-000-121-JIR |
| Papain | Worthington Biochemical | LS003120 |
| PD0332991 isethionate | Sigma Aldrich | PZ0199 |
| Pen-Strep | Sigma-Aldrich | P0781 |
| PhenoPlate™ 96-well microplates | Perkin Elmer | 6055302 |
| PMSF | G-Biosciences | 786-055 |
| Precision Plus Protein Dual Color Standards | BioRad | 1610374 |
| Rock Inhibitor Y-27632 | DNSK International | DNSKKI-15-02 |
| Sera-Mag Beads | Thermo Scientific | 4515-2105-05025,<br>6515-2105-050250 |
| Sequencing grade modified trypsin | Promega | V5111 |
| Sodium ascorbate | Sigma-Aldrich | A7631 |
| Sodium azide | Sigma-Aldrich | S2002 |
| Sodium Bicarbonate | Thermo Fisher | 25080-094 |
| StemFlex Medium | ThermoFisher | A3349401 |
| Streptavidin-Sepharose affinity resin | GE healthcare | 17-5113-01 |
| TMT16pro labelling reagent | Thermo Scientific | A44522 |
| Trolox | Abcam | ab120747 |
| Tryple-Express | Life Technologies | 12604021 |
| **Commercial kits** |  |  |
| Pierce protein assay kit | ThermoFisher Scientific | 22660 |
| **Experimental models: Cell lines** |  |  |
| KOLF2.1J | Human induced pluripotent stem cell line | RRID:CVCL_B5P3 |

### Human induced pluripotent stem cell (hiPSC)-derived hypothalamic neuron culture

KOLF2.1J (P24-P38) (Pantazis et al. 2022) and WTC-11 (P40-P45) (Kreitzer et al. 2013) hiPSC lines were maintained under feeder-free conditions on 0.01% Geltrex-coated plates in StemFlex medium without antibiotics and passaged with EDTA when 70-80% confluent (H.-J. C. Chen et al. 2023). Human iPSCs were differentiated into hypothalamic neurons using small molecule modulators as previously described (H.-J. C. Chen et al. 2023; Kirwan, Jura, and Merkle 2017; Merkle et al. 2015). Dissociated cells were plated onto 0.01% Geltrex-coated plates at a density of 1 × 10^5^ cells/cm^2^ in StemFlex medium supplemented with 10 μg/mL ROCK Inhibitor Y-27632. The following day, medium was changed to N2B27 medium (1:1 Neurobasal-A to DMEM/F12 with GlutaMAX media, 1× Glutamax, 0.075% Sodium bicarbonate, 1× MEM NEAA, 200 mM ascorbic acid, 1× penicillin-streptomycin, 1× B27 supplement, and 1× N2 supplement) and sequentially treated with signaling pathway modulators (H.-J. C. Chen et al. 2023). On day 13, progenitors were dissociated, replated on 4 μg/mL laminin 511-coated plates in N2B27 medium with 10 μg/mL BDNF and 10 μg/mL ROCK inhibitor at a concentration of 3 × 10^5^ cells/cm^2^. The day after replating, media were changed to Synaptojuice medium 1 (N2B27 medium, 2 μM PD0332991 isethionate, 5 μM DAPT, 370 μM CaCl2, 1 μM LM22A4, 2 μM CHIR99021, 300 μM GABA, and 10 μM NKH447) supplemented with 10 μg/mL BDNF. After 1 week, media were switched to Synaptojuice 2 (N2B27 medium, 2 μM PD0332991 isethionate, 370 μM CaCl2, 1 μM LM22A4, and 2 μM CHIR99021) and cells were cultured for at least another week (day 30 onwards) before use in experiments.

### Proximity labelling by APEX2

For proximity labelling using APEX2 (May et al. 2021), 50 mM biotin-phenol (dissolved in 100% DMSO) was added to the medium to a final concentration of 500 μM and incubated for 30 minutes at 37°C. Subsequently, H_2_O_2_ was added to the medium to a final concentration of 1 mM and incubated for 3 minutes at room temperature with gentle shaking to initiate the biotinylation reaction. The reaction was quenched by transferring cells to ice and replacing the medium with ice-cold buffer containing 10 mM sodium ascorbate and 10 mM sodium azide in 1x PBS. Cells were washed two more times with the same buffer supplemented with 5 mM Trolox and the last wash was left for 5 minutes. Cells in 96-well PhenoPlate microplates were then fixed with 4% PFA for 20 minutes at room temperature for immunofluorescence, while cells in 10 cm plates were immediately lysed in 500 µl of 1x RIPY+Q lysis buffer (Table 1) for downstream applications such as western blot, streptavidin chromatography, and mass spectrometry.

**Table 1:**
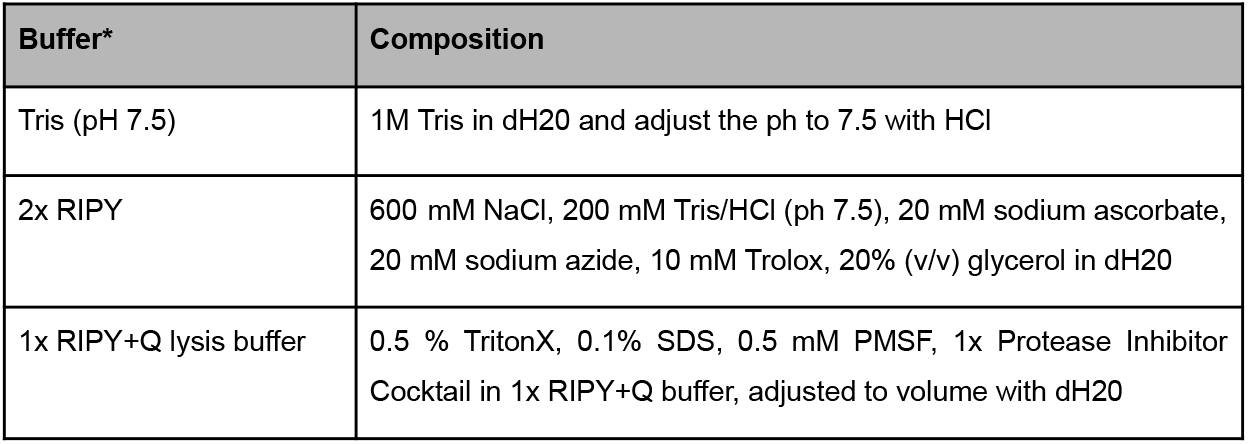
Buffers used for cell lysis. *All buffers were made fresh before use.

### Immunofluorescence and Imaging

Fixed cells on 96-well PhenoPlate microplates were blocked and incubated with primary and secondary antibodies (Table 2), as previously described (Macarelli et al. 2024). For visualisation of biotinylated proteins, cells were incubated overnight at 4 °C in blocking solution and then incubated with 1:1000 488-conjugated Streptavidin for 2 h at RT 1:1000 in blocking solution. Images were captured using a Perkin Elmer Opera Phenix Plus High-Content Screening System capturing images with 10–15 optical planes per Z-stack at 1 μm intervals, processed with ImageJ, and assembled with Affinity Designer. For quantification of cilia length, cells were incubated with 200 nM Janelia Fluor HaloTag Ligand (Grimm et al. 2015) JF503/635i or 1 μM Promega HaloTag Alexa Fluor 488 Ligand and primary cilia length was measured with the previously described pipeline (Macarelli et al. 2024) in Harmony high-content imaging and analysis software. Streptavidin intensity in Scarlet + structures and surrounding regions was quantified using the Harmony software. Briefly, objects in the 555 channel (3xCTS construct) with an area bigger than 4 µm^2^, length smaller than 20 µm, roundness value lower than 0.8, and intensity value higher than 36, were automatically selected. Objects falling on imaging field borders were excluded from analysis. Finally, the mean intensity of the 488 channel (Streptavidin) was calculated in the selected objects and normalised to the mean signal intensity of the adjacent surrounding regions for each object.

**Table 2:** Antibodies used for immunostaining.

| Primary antibodies |  |  |  |
| --- | --- | --- | --- |
| Antibody | Species | Dilution | Catalogue |
| anti-MAP2 | Chicken | 1:2000 | Abcam ab5392 |
| anti-ARL13B | Rabbit | 1:700 | Proteintech 17711-1-AP |
| anti-acetylated tubulin | Mouse | 1:2000 | Sigma T7451 |
| anti-RFP (Scarlet) | Mouse | 1:500 | Synaptic System 409 011 |
| anti-POMC (ACTH/αMSH) | Mouse | 1:5000 | Prof. Anne White |
| anti-LPAR1 | Rabbit | 1:200 | Proteintech 20442-1-AP |
| anti-MYOF | Rabbit | 1:80 | HPA030278 |
| anti-NRP1 | Rabbit | 1:75 | HPA014245 |
| anti-OSBPL10 | Rabbit | 1:110 | HPA003636 |
| anti-PTPRM | Rabbit | 1:110 | HPA003891 |
| anti-GNAI1 | Rabbit | 1:55 | HPA042141 |
| anti-EPB41L3 | Rabbit | 1:57 | HPA028605 |
| Secondary antibodies |  |  |  |
| Antibody | Species | Dilution | Catalogue |
| Alexa anti-rabbit 488 | Donkey | 1:500 | Thermo Fisher Scientific A21206 |
| Alexa anti-mouse 555 | Donkey | 1:500 | Thermo Fisher Scientific A31570 |
| Alexa anti-chicken 647 | Donkey | 1:500 | Strattech 703-606-155 |
| Alexa Fluor™ 488 - Streptavidin | / | 1:1000 | ThermoFisher Scientific S11223 |
| HaloTag Alexa Fluor 488 Ligand | / | 1:1000 | Promega G1002 |
| JF503 HaloTag Ligand | / | 1:5000 | Janelia Materials |
| JF635i HaloTag Ligand | / | 1:17,000 | Janelia Materials |
| **DNA stain** |  |  |  |
| Molecular Probes Hoechst 33342, Trihydrochloride, Trihydrate |  | 1:10,000 | Thermo Fisher Scientific H3570 |
| DAPI (4',6-diamidino-2-phenylindole) |  | 1:1000 | ThermoFisher Scientific D1306 |

### SDS-PAGE and western blotting

After cells were lysed in 1x RIPY+Q lysis buffer (Table 1), lysates were left on ice for 20 minutes and then centrifuged at maximum speed (15.000 rpm) at 4°C for at least 30 minutes (30 min to 1 h). Protein concentration was quantified using the Pierce 660nm Protein Assay Kit. For SDS-PAGE, samples were diluted in 5x Sodium Dodecyl Sulfate (SDS) sample buffer (Table 3) supplemented with 50 mM Dithiothreitol (DTT) and left 20 minutes at room temperature before running on 4-15% polyacrylamide gel at 100V for 15 minutes followed by 130V for 100’. Samples were transferred to nitrocellulose or PVDF membranes and stained for total protein content using LICOR Revert Total Protein Stain. Membranes were then blocked in LICOR TBS blocking solution for 1 hour at room temperature and immunoblotted overnight at 4 °C on an orbital shaker with primary antibodies (Table 4) diluted in 5% milk in TBS and 1 hour at room temperature with secondary antibodies (Table 4). Finally, membranes were washed three times for 5 minutes with 1x TBS-T and imaged using the LICOR Odyssey imaging system. Quantification of western blot signal intensities was performed in a three-step normalization process. First, total protein stain intensities were obtained for each lane and normalized by dividing each value by the highest total protein intensity observed across all samples. Second, the intensity of the target protein in each lane was divided by the corresponding normalized total protein value to account for loading differences. Third, each resulting value was normalized to the corresponding value from the wild-type (WT) control labeled sample, to calculate enrichment of target proteins compared to control.

**Table 3:**
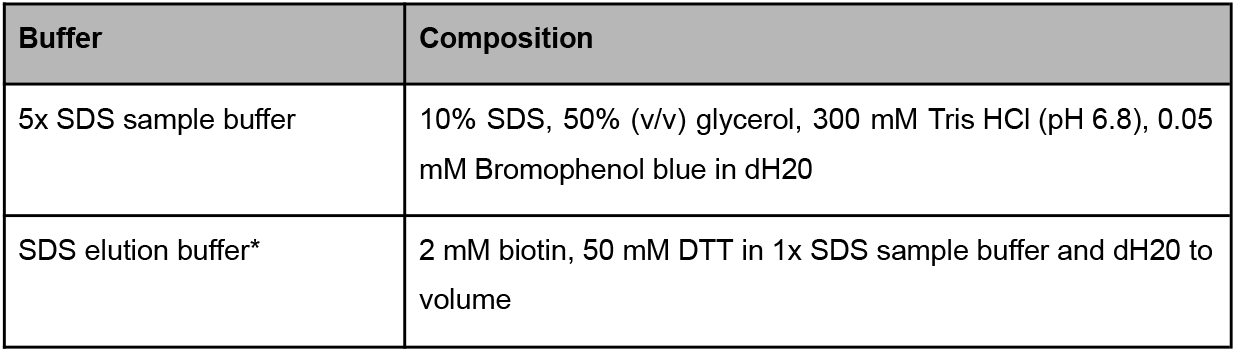
Buffers used for SDS-PAGE. *Add DTT fresh. SDS elution buffer can be stored at -20°C.

| Buffer | Composition |
| --- | --- |
| 5x SDS sample buffer | 10% SDS, 50% (v/v) glycerol, 300 mM Tris HCl (pH 6.8), 0.05 mM Bromophenol blue in dH2O |
| SDS elution buffer* | 2 mM biotin, 50 mM DTT in 1x SDS sample buffer and dH2O to volume |

**Table 4.** Antibodies used for western blotting. M: mouse, rb: rabbit

| Antibody | Species | Dilution | Catalogue |
| --- | --- | --- | --- |
| anti-IFT88 | Rabbit | 1:1000 | 13967-1-AP |
| anti-IFT57 | Rabbit | 1:1000 | 11083-1-AP |
| anti-IFT20 | Rabbit | 1:1000 | 13615-1-1AP |
| anti-RFP (Scarlet) | Mouse | 1:500 | SS 409 011 |
| **Secondary antibody** |  |  |  |
| Streptavidin 680RD | / | 1:1000 | 926-68079 LICORbio |
| anti-rabbit 800CW | Donkey | 1: 10000 | 926-32213 LICORbio |

### Streptavidin chromatography

Following proximity labelling and cell lysis, biotinylated proteins were isolated using streptavidin-sepharose chromatography as previously described (May et al. 2021). Cell lysates containing 20 mg of total protein were diluted to a concentration of 4.4 mg/ml in 4.5 ml lysis buffer were incubated with 200 μl of 50% streptavidin-sepharose bead slurry (100 μl bead bed volume) for 1 hour at room temperature while rolling. Beads had been previously equilibrated by washing once with 1x PBS and twice with 1x RIPY+Q lysis buffer. After incubation, beads were centrifuged, and the unbound material was saved for analysis. The beads were then subjected to 10 washes with 500 μl of lysis buffer and centrifuging at 200 x g for 30 seconds, followed by 5 similar washes with 2 M urea buffer made fresh before use in10 mM Tris pH 7.5.

For western blot analysis, 1/6th of the bead-bound proteins were eluted by boiling in 30 μl of 1x SDS elution buffer containing 2 mM biotin for 5 minutes at 95°C while shaking at 900 rpm, followed by centrifugation at 200 x g for 2 minutes. Eluted and unbound proteins were prepared for western blot analysis as described above and used to assess streptavidin pull-down efficiency prior to MS analysis.

To prepare samples for mass spectrometry (MS) analysis, all equipment and reagents were thoroughly cleaned, buffers were filter-sterilized, MS-grade water was used and all buffers were prepared fresh. After streptavidin chromatography, 5/6th of the bead-bound proteins were subjected to on-bead reduction, alkylation, and digestion with Lys-C for 4 hours at 37°C followed by overnight digestion with trypsin (Wiśniewski et al. 2009; May et al. 2021).

### TMT-based LC-MS/MS proteomics

The EMBL proteomics core facility (Heidelberg, Germany) processed and analyzed samples for protein quantification using mass spectrometry (MS). Peptides were cleaned-up via a reverse-phase chromatography step (OASIS HLB 96-well µElution plate), reconstituted in 10 µl H2O and reacted for 1 h at room temperature with TMT16pro labelling reagent. To this end, 50 µg of TMT16pro label reagent were dissolved in 4 µl of acetonitrile and added to the peptides. Excess TMT reagent was quenched by the addition of 4 µl of an aqueous 5% hydroxylamine solution. Peptides were reconstituted in 0.1 % formic acid, and equal volumes of each uniquely labelled sample were mixed. Mixed peptides were purified by a reverse phase clean-up step (OASIS HLB 96-well µElution Plate). Peptides were subjected to an off-line fractionation under high pH conditions (Hughes et al. 2014) yielding 12 fractions.

Peptides were separated using an Ultimate 3000 nano RSLC system (Dionex) equipped with a trapping cartridge (Precolumn C18 PepMap100, 5 mm, 300 μm i.d., 5 μm, 100 Å) and an analytical column (Acclaim PepMap 100. 75 × 50 cm C18, 3 mm, 100 Å) connected to a nanospray-Flex ion source. The peptides were loaded onto the trap column at 30 µl per min using solvent A (0.1% formic acid) and eluted using a gradient from 2 to 80% Solvent B (0.1% formic acid in acetonitrile) over 2 h at 0.3 µl per min (all solvents were of LC-MS grade). The Orbitrap Fusion Lumos was operated in positive ion mode with a spray voltage of 2.2 kV and capillary temperature of 275 °C. Full scan MS spectra with a mass range of 375–1500 m/z were acquired in profile mode using a resolution of 120,000 with a maximum injection time of 50 ms, AGC operated in standard mode and a RF lens setting of 30%.

Fragmentation was triggered for 3 s cycle time for peptide like features with charge states of 2–7 on the MS scan (data-dependent acquisition). Precursors were isolated using the quadrupole with a window of 0.7 m/z and fragmented with a normalized collision energy of 34%. Fragment mass spectra were acquired in profile mode and a resolution of 30,000. Maximum injection time was set to 94 ms and AGC target to custom. The dynamic exclusion was set to 60 s.

### MS data analysis

Acquired data were analyzed using FragPipe (Kong et al. 2017) and a Uniprot Homo sapiens fasta database (UP000005640, ID9606, 20.594 entries, date: 26.10.2022, downloaded: 11.01.2023) including common contaminants. The following modifications were considered: Carbamidomethyl (C, fixed), TMT10plex (K, fixed), Acetyl (N-term, variable), Oxidation (M, variable) and TMT10plex (N-term, variable). The mass error tolerance for full scan MS spectra was set to 10 ppm and for MS/MS spectra to 0.02 Da. A maximum of 2 missed cleavages were allowed. A minimum of 2 unique peptides with a peptide length of at least seven amino acids and a false discovery rate below 0.01 were required on the peptide and protein level (Savitski et al. 2015).

The raw output files of FragPipe were processed using the R programming environment. 7143 proteins passed the quality control filters. In order to correct for technical variability, batch effects were removed using the ‘removeBatchEffect’ function of the limma package (Ritchie et al. 2015) on the log2 transformed raw TMT reporter ion intensities. Subsequently, normalization was performed using the ‘normalizeVSN’ function of the limma package (VSN - variance stabilization normalization - (Huber et al. 2002)). Normalization was performed separately for the following groups: minus and plus labelling. This ensured that abundance differences across these groups were preserved. Missing values were imputed with the ‘knn’ method using the ‘impute’ function from the Msnbase package (Gatto and Lilley 2012). This method estimates missing data points based on similarity to neighboring data points, ensuring that incomplete data did not distort the analysis. Differential expression analysis was performed using the moderated t-test provided by the limma package (Ritchie et al. 2015). The model accounted for replicate information by including it as a factor in the design matrix passed to the ‘lmFit’ function. Imputed values were assigned a weight of 0.01 in the model, while quantified values were given a weight of 1, ensuring that the statistical analysis reflected the uncertainty in imputed data. Proteins were annotated as hits if they had a false discovery rate (FDR) below 0.05 and an absolute fold change greater than 2. Proteins were considered candidates if they had an FDR below 0.2 and an absolute fold change greater than 1.5.

Metascape (metascape.org) was used for GO term enrichment analysis (Fisher’s exact test <0.05, ≥3 genes, enrichment factor >1.5) across biological processes (BP), molecular functions (MF), and cellular components (CC). Enriched terms were clustered based on similarity and analysis type (BP, CC and MF), with the most enriched term representing each cluster. The full MS gene set served as the background for enrichment. Active subnetworks were identified using DOMINO, a computational tool that integrates a list of input proteins with a reference protein-protein interaction network (Levi, Elkon, and Shamir 2021) such as STRING.

## Statistical analysis

All data plotted and analysed using GraphPad Prism (Version 10.2.0) were evaluated for normality using a Shapiro-Wilk test and then analysed using one- or two-way analysis of variance (ANOVA) followed by Bonferroni post hoc test, or Kruskal–Wallis test followed by Dunn’s post hoc test when comparing more than two datasets. Mann–Whitney test was used to compare two datasets. p values < 0.05 were considered to be statistically significant. All frequency distributions were generated using relative frequency (percentages) and all bins width were setted to 1.5. Data are presented as mean ± standard error of the mean (SEM) or individual values ± SEM, and statistical tests used are indicated in the figure legends.

## Results

### APEX2 targeting to the cilium and specific biotinylation of ciliary proteins of hiPSC-derived hypothalamic neurons

In order to characterise the proteome of primary cilia of human iPSC-derived hypothalamic neurons, we employed APEX2-based proximity labelling to selectively label and isolate ciliary proteins. To allow proximity labelling, we cloned the peroxidase APEX2 into transmembrane constructs with identical components except for the absence or presence of a short ciliary targeting sequence (CTS) from the MCHR1 protein to target the construct to the plasma membrane (TM) or primary cilia (3xCTS) compartments (Macarelli et al. 2024). Both constructs contain an extracellular HaloTag, a trans-membrane domain (TMD), the peroxidase APEX2 and mScarlet-I fluorescent protein, and homology arms that facilitate targeted integration into the human *AAVS1* safe-harbor locus (Figure 1A). When incubated with hydrogen peroxide (H_2_O_2_), APEX2 converts exogenously supplied biotin-phenol (BP) to biotin-phenoxyl radicals which can label proteins in a small radius (20 nm) (Rhee et al. 2013). Biotinylated proteins can then be extracted from cell lysates and isolated through a chromatography column where they bind to streptavidin-conjugated beads to enable elution and analysis by western blot or mass-spectrometry (MS) (Figure 1B). We targeted TM and 3xCTS constructs to the *AAVS1* locus of the KOLF2.1J iPSC line to generate stable cell clones that we differentiated in parallel to hypothalamic neurons. We then biotinylated these cultures alongside control samples generated from wild-type (WT) KOLF2.1J-derived hypothalamic neurons, and mock-labeled (H_2_O_2_ omitted) controls (Figure 1B).

**Figure 1:**
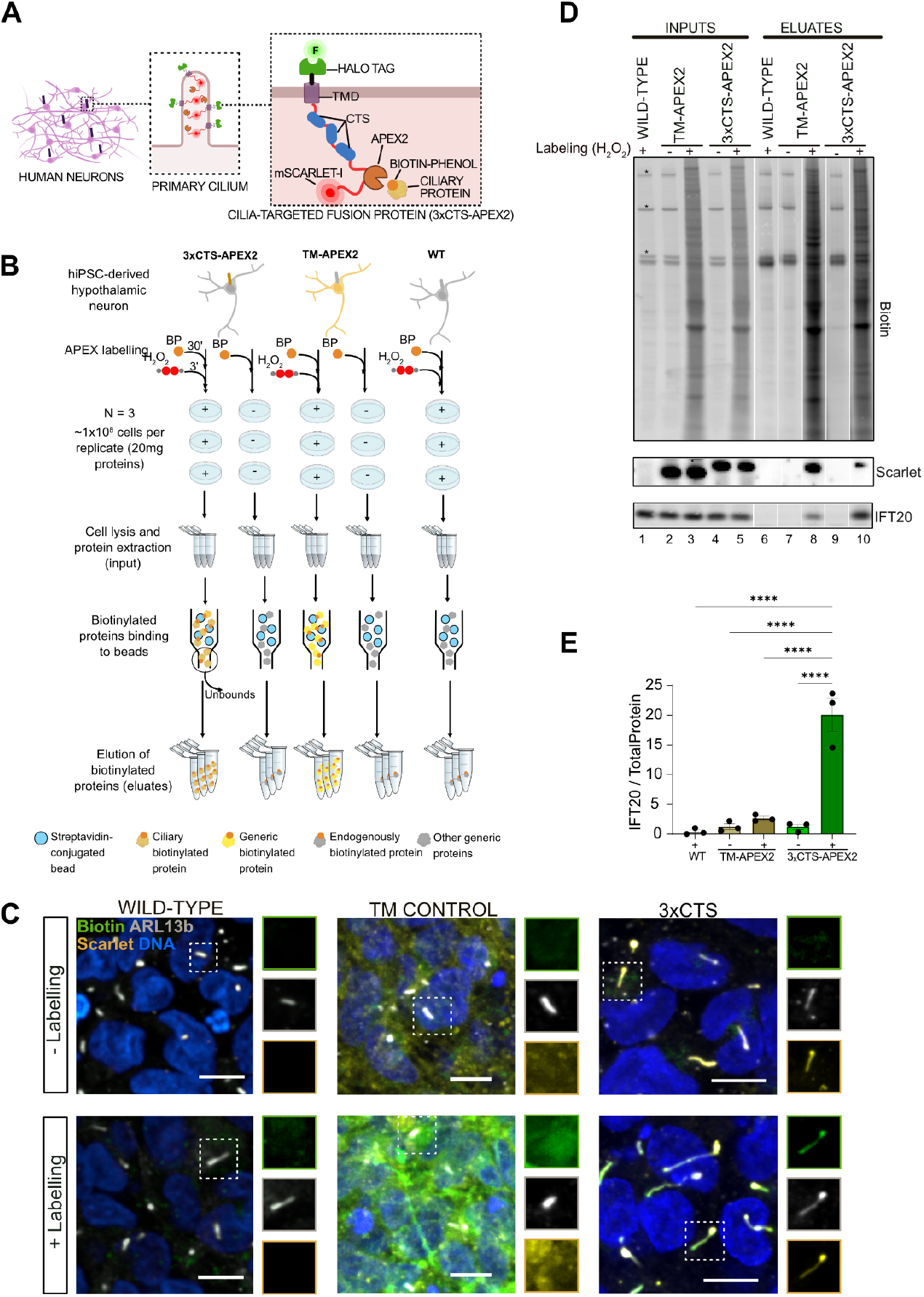
APEX2-mediated biotinylation of ciliary proteins of hypothalamic neurons. **(A)** Schematic of the 3xCTS construct modified to carry APEX2 and stably integrated in hiPSCs, which express the reporter in primary cilia when differentiated to hypothalamic neurons. **(B)** Hypothalamic neurons derived from 3xCTS-APEX2, TM-APEX2 and WT hiPSC lines were incubated with BP (biotin-phenol) for 30’ and H_2_O_2_ (hydrogen peroxide) for 3’ to induce APEX2-mediated biotinylation. Around 1×10^8^ cells per replicate (N = 3) were lysed to extract 20 mg of proteins each, 0.1% of these were used as “input” in western blot. Biotinylated proteins were then run through a chromatography column where they bound to streptavidin-conjugated beads, and material that did not bind to the beads was collected (unbounds). Each replicate was run independently. After washing, proteins were eluted from the beads and either further processes for mass spectrometry analysis or used for western blot. **(C)** Representative images showing 3xCTS-APEX2, TM-APEX2 and WT hypothalamic neurons in the absence (−labelling) or presence (+ labelling) of labelling reagents. Cilia are identified as ARL13B+ structure (grey), the fusion construct is visualised using Scarlet (yellow) and biotin using streptavidin (green). Nuclei are in blue. Scale bars represent 10 μm, boxes show channel-splitted cilia. MAP2: neuronal marker; POMC: hypothalamic POMC neurons marker. **(D)** Western blot (WB) analysis of proteins extracted from 3xCTS-APEX2, TM-APEX2 or WT hypothalamic neurons after APEX2-mediated labelling (+) or mock labelling (−). Proteins were collected immediately after cell lysis (inputs) or after purification by streptavidin chromatography (eluates). Biotinylated proteins were detected by streptavidin-HRP, while other proteins were detected by specific antibodies. Asterisks indicate endogenous biotinylated proteins, samples were run in triplicates but only one representative is shown. Uncut blots are shown in supplementary figures. **(E)** Quantifications of the signal intensity of IFT20 normalised to total protein stain in the eluate samples. **** p < 0.0001 by Kruskal–Wallis test.

To assess the specificity of ciliary labelling, we stained cultures with Alexa Fluor 488-labeled streptavidin. Incubation with both biotin-phenol and H_2_O_2_ (the ‘+ labelling’ condition) induced specific labelling of primary cilia in the 3xCTS neurons, non-ciliary labelling in TM control neurons, and only background labelling in WT control neurons (Figure 1C). We optimised APEX2-mediated biotinylation of ciliary proteins by testing some of the most commonly used incubation times for biotin-phenol (45’ and 30’) in the literature (Hung et al. 2016; Mick et al. 2015; May et al. 2021) and four different hydrogen peroxide incubation times (0’,1’,2’,3’), and selected the condition showing the best noise-to-signal ratio (30’ BP and 3’ H_2_O_2_) for subsequent studies (Figure S1A-D). We confirmed biotinylation of ciliary proteins in neurons positive for neuronal (MAP2) and hypothalamic (POMC) markers (Figure S1E). We also found that 53.1% ± 1.78 of DAPI+ cells expressed the cilia-targeted construct (Scarlet +) (Figure S1F), which would represent the population of ciliated cells as we previously showed complete colocalization (>99%) of our construct with the ciliary protein ARL13B (Macarelli et al. 2024). By western blots, we confirmed the efficient biotinylation of proteins derived from 3xCTS and TM cultures (Figure 1D), and the enrichment of cilia-associated proteins such as IFT20 by pulldown (Figure 1D,E), but not of IFT88 and IFT57 (Figure S1G-I and Figure S2). Together, these findings demonstrate the selective labelling of primary cilia in neurons derived from the stable KOLF2.1J-3xCTS cell line.

### Mass spectrometry analysis of ciliary proteins of hypothalamic neurons reveals proteins involved in synapses assembly

We subjected 3xCTS, TM control and WT control labelled and mock-labelled lysates in triplicate to Tandem mass tag (TMT)- based liquid chromatography-mass spectrometry (LC-MS/MS). Triplicate samples showed high reproducibility, and known ciliary proteins, such as ARL13B and Smoothened (SMO), were found enriched in 3xCTS labelled samples (MCH_plus) compared to TM control (TM_plus) and WT (WT_plus) (Figure S3A,B). Very little signal of biotinylated proteins was detected from WT and mock-labelled controls (Figure S3C,D), so to ensure a more even comparison we compared the abundance of biotinylated proteins pulled down in cilia-labelled samples (3xCTS) to those pulled down in labelled control (TM) samples (Figure 2A). Only proteins quantified with at least two unique peptide matches and in at least 2 replicates were considered for the analysis. We then defined ‘hit’ proteins as enriched at least 2-fold in 3xCTS samples with an FDR<0.05, and ‘candidate’ proteins as enriched at least 1.5-fold with an FDR<0.2, and identified 141 hit and 408 candidate proteins in neuronal primary cilia. The hits included the APEX2 construct itself and well-known-ciliary proteins including ARL13B, SMO, TUB, GLI3, CFAP20, CLUAP1, INPP5E, and DCDC2, and components of the IFT machinery such as IFT81, IFT57, IFT122 and IFT172 were present among the candidates (Figure 2A).

**Figure 2:**
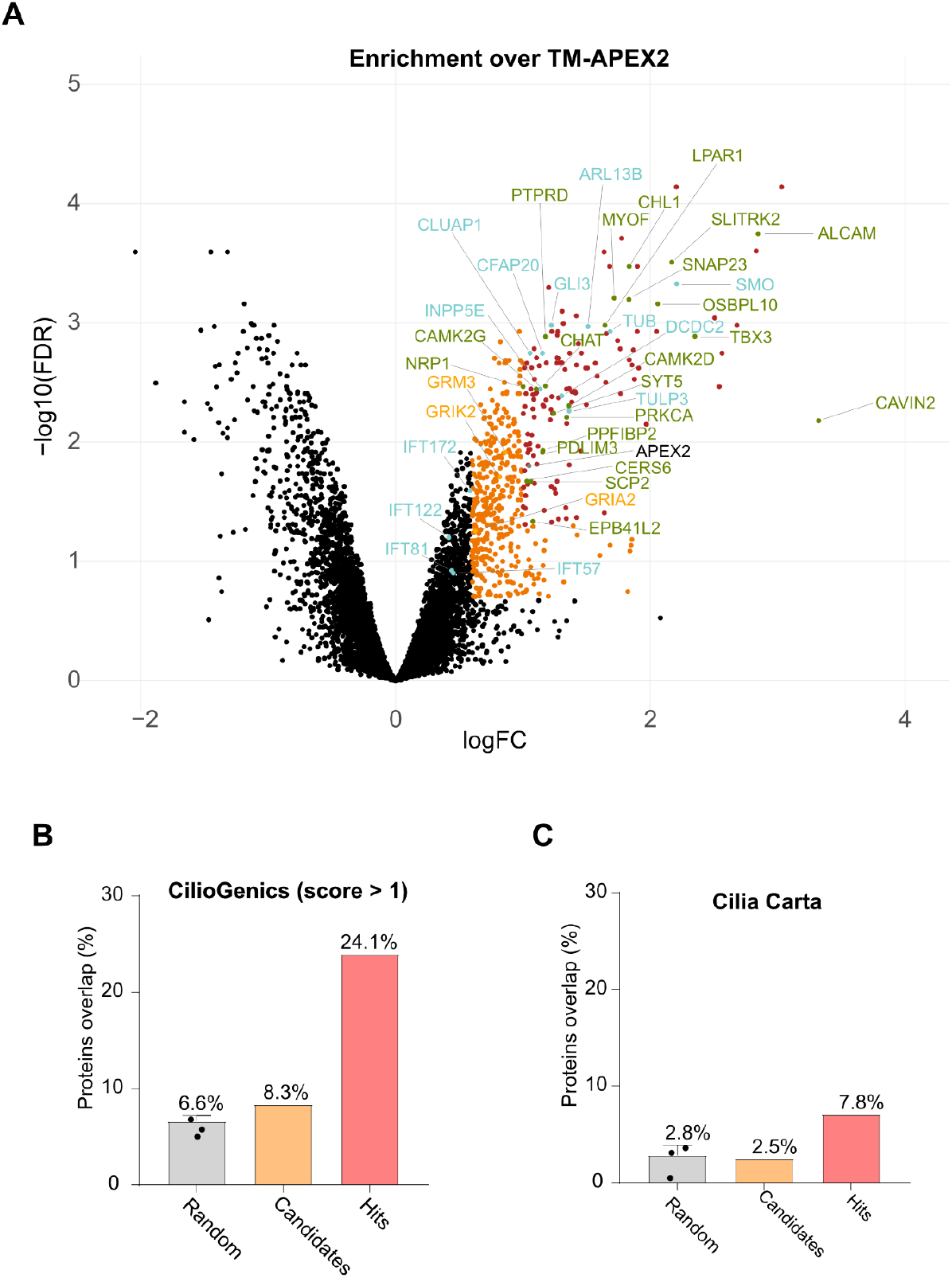
Proteomics of primary cilia of hiPSC-derived hypothalamic neurons. **(A)** Volcano plots of statistical significance (FDR) versus protein enrichment (FC) in 3xCTS-APEX2 labelled (plus) compared to labelled (plus) TM-APEX2 control, using limma analysis. Proteins categorised as hits (FDR ≤ 0.05 and FC ≥ 2) are in red, as candidates (FDR ≤ 0.2 and FC ≥ 1.5) are in orange, some known ciliary proteins are in blue and some hits and candidates further analysed below are in green. **(B)** The lists of hit and candidate proteins and 3 lists of random human genes were compared to proteins predicted to be ciliary in the CilioGenic database (score >1, high probability). **(C)** As in B but compared to the Cilia Carta database. Bars represent mean + SEM, 6.6% ± 0.6 in B and 2.8% ± 1 in C.

We further confirmed the enrichment of ciliary proteins in our dataset by comparing the list of candidate and hit proteins to similarly-sized lists of randomly selected genes or ciliary genes as represented in the CilioGenics database (Pir et al. 2024) and Cilia Carta database (van Dam et al. 2019) and observed a substantial enrichment in ciliary proteins among hits (Figure 2B,C). To confirm and extend this analysis, we compared cilia-enriched hit and candidate genes to other published ciliary databases derived mostly from mouse models but also from human cell lines, and curated lists (Sigg et al. 2017; Y.-D. Chen et al. 2023; Vasquez, van Dam, and Wheway 2021; Turan et al. 2023; Pir et al. 2024; van Dam et al. 2019; Nevers et al. 2017; Mick et al. 2015; Gheiratmand et al. 2019; Gupta et al. 2015; Kohli et al. 2017) (Figure S3E). We found that approximately 16% of cilia-enriched proteins had been previously mapped to the centrosome-cilium interface (Gupta et al. 2015), suggesting biotinylation of proteins in the 3xCTS cell line at the ciliary base as well as the main structure.

To gain functional insights into the identified hit proteins, we carried out gene ontology (GO) analysis (Carbon et al. 2021) using Metascape (Zhou et al. 2019) and considering biological processes (BP), molecular functions (MF), and cellular components (CC). A total of 261 terms (fisher’s exact test value < 0.05, minimum count of 3 genes, and enrichment factor > 1.5) were grouped into clusters based on their membership similarities and GO term analysis type (BP, MF or CC), with the most enriched term chosen to represent the cluster (Figure 3 A,B and S4A,B). Enriched functions were mainly related to cell-cell contact, synapse and neurotransmitter function, axon guidance, and sterol binding. To test for potential interaction between cilia-enriched proteins, we used the DOMINO (Levi, Elkon, and Shamir 2021) tool on STRING protein-protein interaction data (Szklarczyk et al. 2023) to identify modules of interacting proteins. We named modules based on their most enriched GO term, and ranked modules according to their STRING clustering score. Hit ciliary proteins mainly scored in the ‘axon guidance’ and ‘synapse assembly modules, including proteins like NRP1, CHL1, TBX3, PDLIM3, PTPRD, and SLITRK2 (Figure 3C,D). These networks also contained candidate ciliary proteins including the glutamate receptors GRM3, GRIA2, GRIK2, and GRID1. Other modules containing candidate ciliary proteins included proteins functioning in the GABAergic synapse (Figure 3E), whereas proteins depleted in cilia (FDR ≤ 0.05 and FC ≤ - 2) had roles in DNA replication (Figure S4C).

**Figure 3:**
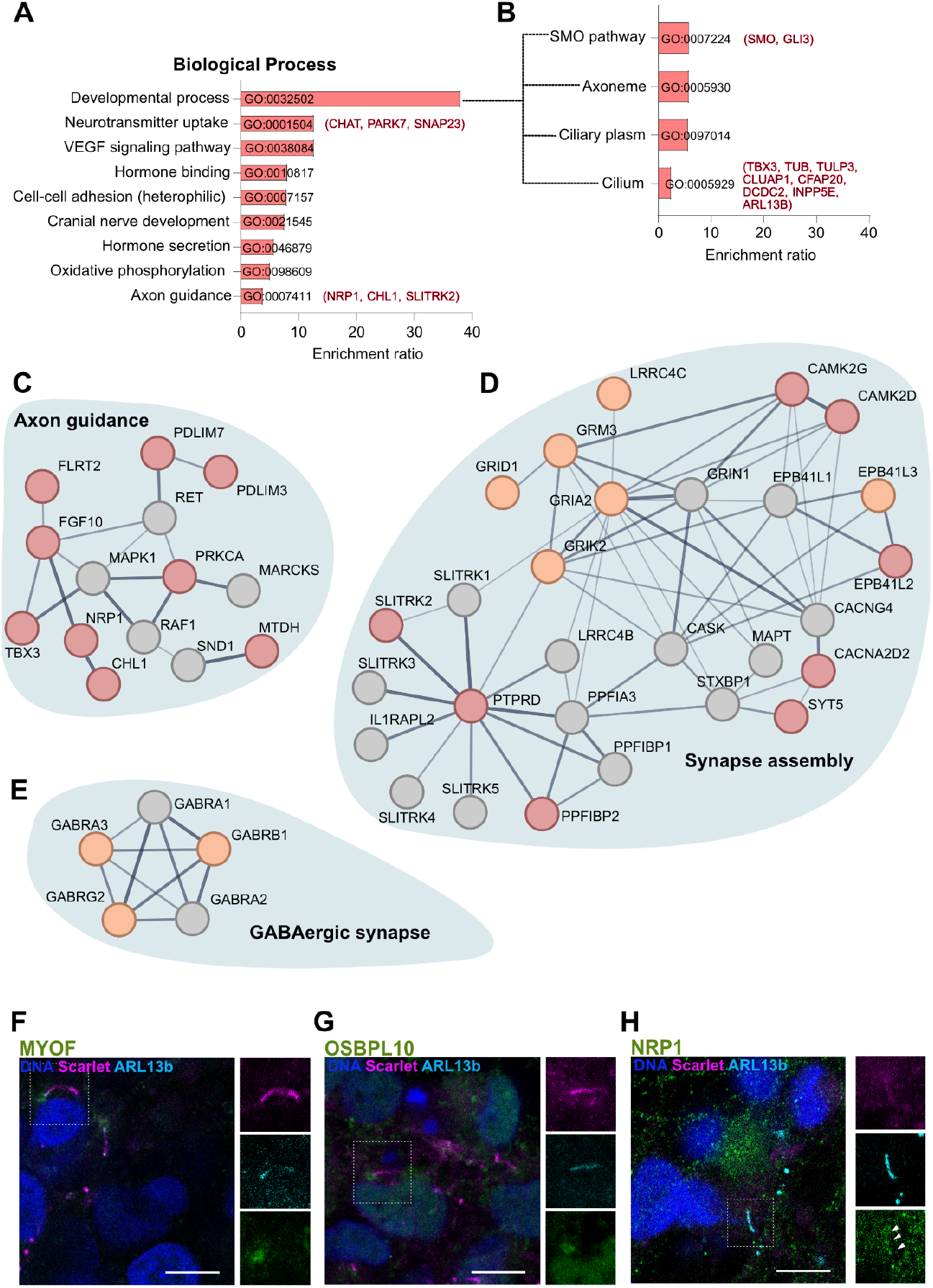
Proteins identified by ciliary proteomics in hypothalamic neurons are involved in axon guidance and synapses. **(A,B)** GO term enrichment analysis of hit proteins. Each module represents the most enriched term (fisher’s exact test value < 0.05) in a cluster of terms with membership similarities. (B) The ‘developmental process’ module contains several clia-related terms. Between brackets some example proteins of the module. **(C-E)** Active networks identified using DOMINO. Red and orange nodes identify hit and candidate proteins respectively, grey nodes indicate proteins either not detected or not enriched in the dataset. The edge thickness is determined by the STRING level of confidence of the data. Module names indicate the most enriched GO term (C,D). The first two highest scoring modules are shown for the list of hits and (E) the first highest scoring module for the list of candidates. **(F-H)** Immunofluorescence staining for some of the hits in hiPSC-derived hypothalamic neurons showing localisation to the cilium base (MYOF, OSBPL10) or to the cilium (NRP1). Scale bars represent 10 μm, insets show channel-splitted cilia. MYOF: myoferlin; NRP1: neuropilin 1; OSBPL10: Oxysterol Binding Protein Like 10.

To further confirm our results, we performed immunofluorescence stainings with antibodies from the Human Protein Atlas (Thul et al. 2017; Hansen et al. 2024) to confirm the subcellular localisation of hit and candidate ciliary proteins (Figure 3 F-H and S4D). We found that myoferlin (MYOF) and Oxysterol Binding Protein Like 10 (OSBPL10) were detected at the basal body and that NRP1 was localised along the ciliary shaft. Together these findings confirm and extend previous proteomic studies (Loukil et al. 2023) to human neurons, and suggest functional roles for cilia in synaptic interaction and neurotransmitter sensing.

### The length of hypothalamic primary cilia is regulated by LPA

In the hypothalamus, cilia are likely to be important for metabolic sensing and the regulation of appetite (Brewer et al. 2022) and ciliary localization of Mc4r exhibits dynamic regulation in response to feeding and fasting states in the PVN (Ojeda-Naharros et al. 2025). In addition, primary cilia of the ARC nucleus of the hypothalamus are required for normal body weight regulation (Davenport et al. 2007; D.-F. Guo et al. 2019), but the exact mechanisms behind their role in the ARC nucleus is still unclear. To uncover new links between primary cilia and obesity, we employed GeneCards (Stelzer et al. 2016) to assemble a list of protein-coding genes associated with this disease. We then cross-referenced this list and a list of genes associated with neurological disorders and ciliopathies with our hit list (Figure 4A) and found 84 proteins associated with obesity, of which 57 were also associated with other general neurological disorders and 27 were associated with ciliopathies, including ROR2, TMED7, LRP1B, CFAP20, TUB, ATP5MG, PDLIM7, BAD, and TULP3.

**Figure 4:**
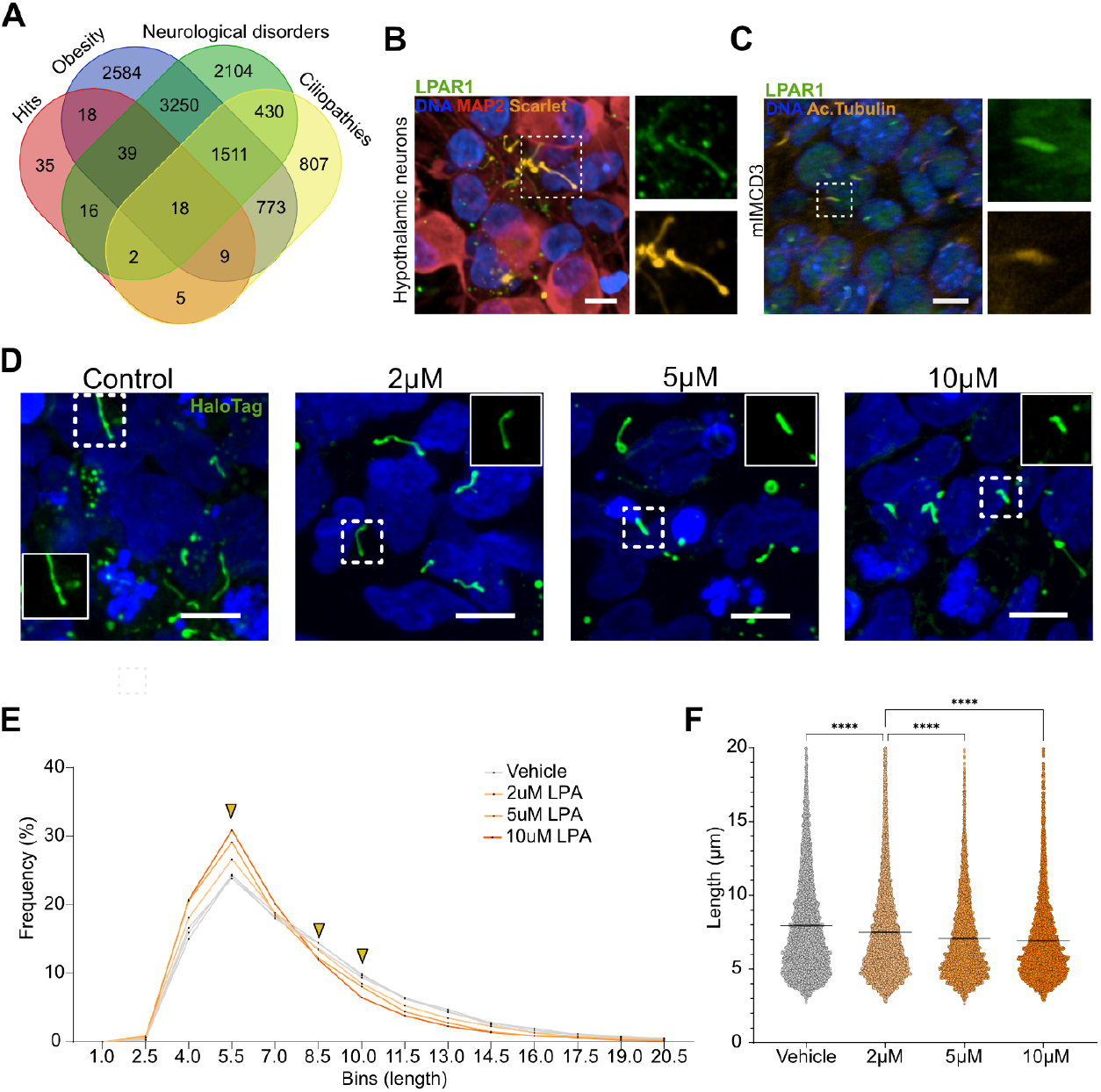
LPA regulates the length of primary cilia of hypothalamic neurons. **(A)** Venn diagram showing the overlap between proteins found in our dataset and proteins associated with neurological disorders, obesity or ciliopathies. **(B)** Immunostaining for LPAR1 in hiPSC-derived hypothalamic neurons expressing the 3xCTS-APEX2 construct (Scarlet). MAP2: neuronal marker. **(C)** Immunostaining for LPAR1 in IMCD3 cells. Scale bars represent 10μm, insets show channel-splitted cilia. Acetylated tubulin: cilia marker; LPAR1: Lysophosphatidic acid receptor 1. **(D)** Representative images of HaloTag+cilia (green) treated with different concentrations of LPA or vehicle (PBS). **(E)** Distribution of the primary cilium length in vehicle and LPA-treated hypothalamic neurons. Cilia from 3 replicates for each condition were pooled together prior to analysing the distribution of lengths. **(F)** Distribution of lengths where each point represents a single cilium and replicates are pooled together. **** p < 0.0001 by Kruskal–Wallis test.

Among the obesity-associated hits, we found that the Lysophosphatidic Acid Receptor 1 (LPAR1) was significantly (3-fold) enriched in cilia. LPAR1 is a member of a group of 6 GPCRs which can bind to lysophosphatidic acid (Geraldo et al. 2021), a bioactive lipid implicated in metabolic disorders (D’Souza, Paramel, and Kienesberger 2018). LPAR1 is the most abundant LPA receptor isoform in the brain, where it plays a role in NPC differentiation and neuronal morphology, motility and excitability (Xiao et al. 2021). Additionally, LPAR1 localises to primary cilia of human astrocytes (Loskutov et al. 2018) and regulates cilia length of mouse NPCs (Hu et al. 2021) and hTERT-immortalized retinal pigment epithelial (RPE-1) cells (Walia et al. 2019). We immunostained iPSC-derived human hypothalamic neurons and the mouse inner medullary collecting duct (mIMCD3) cell line and found that the receptor preferentially localised to primary cilia in both of these cell populations (Figure 4 B,C). To test if LPA can modulate the length of neuronal primary cilia, we treated hiPSC-derived hypothalamic neurons with different concentrations of LPA (2, 5 and 10 μM in PBS) for either 2, 4 or 24h (Figure S5) and quantified primary cilia length by semi-automated analysis of confocal images as previously described (Macarelli et al. 2024). We observed a dose-dependent decrease in cilia length after 24h of LPA treatment (Figure 4D), leading to significant increases in the percentage of shorter cilia (∼ 5.5 μm) and a corresponding decrease in the percentage of longer cilia (∼ 8.5 μm and longer) (Figure 4E,F). These data suggest a functional role for LPA in the regulation of primary cilia length of hypothalamic neurons.

## Discussion

To gain insight into the functional role of primary cilia in human hypothalamic neurons, we employed APEX2-based proximity labelling in an hiPSC line expressing a cilia-targeted reporter (Macarelli et al. 2024). We confirmed this approach selectively labels neuronal primary cilia and enriches for ciliary proteins such as IFT20 (Follit et al. 2006). The recovery of other IFT proteins in control samples might be due to their presence in the Golgi and their accumulation at the ciliary base (Hibbard, Vázquez, and Wallingford 2022). The ∼4-fold enrichment for human ciliary proteins as defined by CilioGenic database may under-represent true enrichment since known hypothalamic ciliary proteins including MCHR1 and SSTR3 are classified as “low probability” in CilioGenic, likely due to the different cell types used to build ciliary protein databases. We also observed enrichment of hedgehog (HH) pathway components (e.g. SMO), which could be attributed to persistent HH signalling in the adult hypothalamus and in our iPSC-derived hypothalamic culture system (Merkle et al. 2015; Antonellis et al. 2021). Finally, the enrichment of lipid-binding proteins (CERS6, SCP2 and OSBPL10) aligns with known functions of lipids in ciliary HH signalling (Nguyen, Truong, and Reiter 2022; Raleigh et al. 2018), and might point to a similar role of other GPCRs. Our approach did not recover many enriched ciliary GPCRs such as MC4R, likely due to both the low abundance of GPCRs and the bias in our differentiation protocol to ARC and ventromedial hypothalamic cell populations (H.-J. C. Chen et al. 2023; Saper and Lowell 2014).

### Limitations of the study

While our study provides the first description of ciliary proteins in human neurons, we acknowledge some intrinsic limitations in our approaches.

TMT-based quantitative proteomics can have lower sensitivity than label-free quantitation methods, limiting power to detect low-abundance proteins. While streptavidin labelling studies suggested that our targeting construct accumulated selectively in primary cilia, it would also have been present in the ER and Golgi, and likely also in the lysosome, leading to the biotinylation of non-ciliary proteins in addition to targets at the cilium. We speculate that the unexpectedly high fraction of basal body- and centrosome-associated proteins we identified might represent construct accumulation at the ciliary base due high expression levels, and that the use of weaker promoters to drive construct expression might lead to greater specificity of subcellular localisation and therefore greater ciliary enrichment in future studies. However, we also note that the comparison was primarily made using published databases generated from mouse cell lines (Mick et al. 2015; Ishikawa et al. 2012; Kohli et al. 2017), whereas the basal body- and centrosome-associated datasets were derived from human cell lines (Gheiratmand et al. 2019; Gupta et al. 2015), which might explain the higher overlap with our dataset.

Endogenous peroxidases can lead to background biotinylation with biotin phenol in the presence of endogenous H_2_O_2_, and this limitation could be reduced by recently-developed methods like iAPEX (Sroka et al. 2025). Protocols leading to higher levels of ciliation could reduce background from construct localisation to non-ciliary compartments (Fig. 1D) though the use of controls such as the isogenic TM cell line should address most of these issues (Aslanyan et al. 2023; Mick et al. 2015; May et al. 2021). While there is always room for improvement, we believe that the dataset and methods we report here are valuable resources for discovering novel human ciliary genes and potential therapeutic targets for obesity such as LPAR1. Moreover, the enrichment of proteins involved in synaptic function and axon guidance provides fresh perspectives on the roles of neuronal cilia in synaptogenesis and neuronal development.

### A potential role for primary cilia in neuronal synapse establishment

We identified 141 potential ciliary proteins, 98 of which were not previously associated with primary cilia. Gene ontology analysis revealed that many of these novel hits are related to synaptic function, such as synapse establishment, neurotransmitter regulation, and cell adhesion. For example, we observed enrichment in choline acetyltransferase (CHAT) which is responsible for the synthesis of acetylcholine (Oda 1999) and multiple proteins involved in membrane trafficking and vesicle release at the synapse (MYOF, SNAP23, SYT5, EPB41L2, CAVIN2). MYOF, a Ca^2+^-triggered vesicle fusion protein (H.-J. C. Chen et al. 2023; Leclère and Dulon 2023), and EPB41L2, erythrocyte membrane protein band 4.1 like 2, have previously been reported as localised to cilia (Aslanyan et al. 2023; Ishikawa et al. 2012; Karunakaran et al. 2020). These findings align with the intriguing possibility that neuronal cilia may fine-tune synaptic activity by sensing neurotransmitter spillover (C. M. Ott et al. 2024) or even by releasing vesicles themselves. Proteins functioning in the GABAergic and glutamatergic synapse were found in mouse neuronal cilia (Loukil et al. 2023) and replicated in our list of candidate proteins, suggesting that neuronal primary cilia might be involved in the regulation of neuronal excitability or gene expression. The ciliary enrichment of Ca^2+^/calmodulin-dependent kinases (CAMK2G and CAMK2D) suggests that neuronal cilia have active calcium signalling downstream of these and other receptors, a finding consistent with observations by others (Hansen et al. 2024) who reported the localization of Ca^2+^/calmodulin-dependent kinases and their regulators to cilia of diverse cell types.

The presence of cell adhesion molecules such as ALCAM, PDLIM3, PTPRD and the liprin protein PPFIBP2, suggests that primary cilia may form physical association with neuronal processes (Wu et al. 2024; Sheu et al. 2022) and with other cilia (Jackson 2012; C. Ott et al. 2012) reminiscent of immune synapses (Stinchcombe et al. 2015; Douanne, Stinchcombe, and Griffiths 2021; Loukil et al. 2023). PDLIM3 has already been found in cilia of fibroblasts where it supports ciliogenesis (Zhang et al. 2023), while ALCAM has been associated with the HH signalling pathway in some cancers (Zhang et al. 2023; G. Chen et al. 2018). Liprins, a family of scaffold proteins with a role in synaptic structure formation, have also been linked to trafficking of GLI proteins to the ciliary tip (Pehkonen, de Curtis, and Monni 2021).

We observed cilia-enriched proteins involved in axon guidance, including NRP1, SLITRK2 and CHL1 and development (TBX3), likely reflecting the cellular heterogeneity in our cultures. Indeed, our differentiation protocol inevitably generates neuronal progenitors and immature cells alongside mature neurons, each potentially contributing distinct ciliary protein signatures. Although improved differentiation protocols might be needed to gain a higher level of homogeneity in the population and cell-type-specific proteome, using hiPSCs allows to generate large numbers of differently derived cell types, as well as patient-derived models. NRP1 has been found in cilia of fibroblasts where it promotes HH signalling (Merkle et al. 2015; Pinskey et al. 2017), while SLITRK2 plays a role in synaptic adhesion (El Chehadeh et al. 2022) SLITRK5 has been found in cilia of osteoblasts where it acts as a negative regulator of HH signalling (El Chehadeh et al. 2022; Sun et al. 2021). Similarly, the transcription factor TBX3 has been shown to regulate GLI3 stability in primary cilia (Emechebe et al. 2016). Taken together, these findings corroborate the established role of cilia in axon development and axonal pathfinding during neural development (Coschiera et al. 2024; Dumoulin et al. 2024; J. Guo et al. 2019; Kumamoto et al. 2012) and validates our approach.

### Regulation of primary cilia length by LPA

We identified LPAR1 as an enriched GPCR in hypothalamic neuronal cilia. LPAR1 and autotaxin (ATX/ENPP2), one of the enzymes responsible for LPA production, are genetically associated with obesity (Birgbauer 2021). Circulating LPA levels are also elevated in obesity and influenced by feeding state (D’Souza, Paramel, and Kienesberger 2018; Dusaulcy et al. 2011; Michalczyk et al. 2017). Given the link between hypothalamic cilia length, sensitivity to metabolic signals, and food intake (Han et al. 2014; Oya et al. 2024), we speculate that LPAR1 may regulate the sensitivity of cilia-mediated metabolic sensing in the hypothalamus (Macarelli, Leventea, and Merkle 2023). LPAR1 and its ligand LPA have been implicated in cilia disassembly via AURKA activation (Hu et al. 2021; Kim et al. 2015) and inhibition of cilia assembly through the PI3K/Akt pathway (Walia et al. 2019). Furthermore, LPA signalling through LPAR1 plays a role in synaptic transmission, mainly in glutamatergic and GABAergic synapses (Peñalver et al. 2017; Roza et al. 2019). After confirming the ciliary localization of LPAR1, we found that the primary cilia of hiPSC-derived hypothalamic neurons where significantly shortened in response to concentrations of LPA previously reported in the literature (Hu et al. 2021; Kim et al. 2015; Walia et al. 2019) and similar to those observed in human plasma and serum (1-15 μM) (Dusaulcy et al. 2011; Michalczyk et al. 2017). The leaky blood-brain-barrier may give nearby hypothalamic neurons access to these LPA concentrations whereas in deeper brain structures LPA concentrations are much lower (0.025–0.2 pM in the cerebrospinal fluid) (Birgbauer 2021; Yung, Stoddard, and Chun 2014). In future studies, it would be interesting to test how LPA treatment alters ciliary receptor composition, signal transduction, and neuronal activity.

## Acknowledgements

We thank Anne White for providing anti-α-MSH antibody, and Dr. Frank Stein and Dr. Per Haberkant at the EMBL proteomics core facility (Heidelberg, Germany) for processing and analysing mass spectrometry samples. We thank Chris Smith of the IMS Tissue and Cell Imaging core facility which is funded by the UKRI Medical Research Council (MRC) Metabolic Disease Unit (MRC_MC_UU_00014/5) and a Wellcome Trust Major Award (208363/Z/17/Z). F.T.M. is a New York Stem Cell Foundation (NYSCF) - Robertson Investigator [NYSCF-R-156], Chan Zuckerberg Initiative (CZI) - Ben Barres Early Career Investigator [CZI NDCN 191942, 10.37921/429861umrcjh], and Wellcome Trust - Sir Henry Dale Fellow [211221/Z/18/Z] and whose funding helped support these studies. V.M. was supported by the Sackler Trust, the MRC Doctoral Training Programme, and the School of Clinical Medicine Cambridge Trust Scholarship. We support inclusive, diverse, and equitable conduct of research. T.J.S. was supported by Deutsche Forschungsgemeinschaft (DFG) funding (TRR152-P28 – Project-ID 239283807). JNH was supported by a Postdoctoral Fellowship from the Wenner-Gren Foundations, and by an EMBO Postdoctoral Fellowship (ALTF 556-2022). For open access, the authors have applied a CC-BY public copyright license to any Author Accepted Manuscript version arising from this submission.

## Notes

### Competing Interest Statement

The authors have declared no competing interest.

