## Supplementary figures for "Proximity proteomics of primary cilia in human hypothalamic neurons"

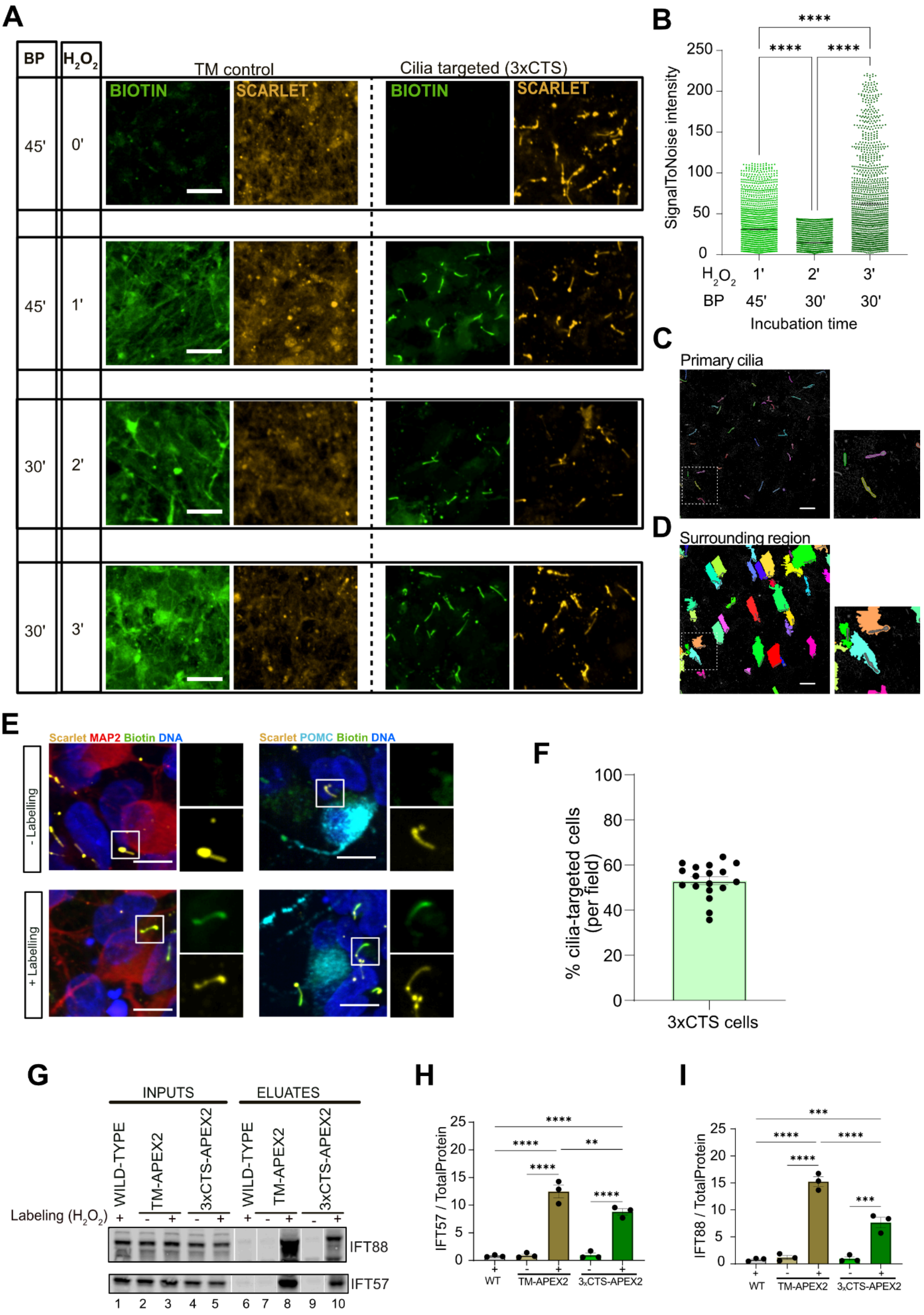

**Figure S1.** **(A)** Immunofluorescence of hiPSC-derived neurons stably carrying either the 3xCTS-APEX2 construct or the TM-APEX2 control construct. Neurons were exposed to biotin-phenol (BP) for 30 or 45 minutes and hydrogen peroxide ($H_2O_2$) for 0,1, 2, or 3 minutes. Expression of fusion constructs is visualised by Scarlet fluorescence (yellow) and biotinylation by streptavidin fluorescence (green). Scale bars represent 10 µm. **(B)** Quantification of streptavidin mean signal intensity in Scarlet+ structures normalised to the mean intensity of the surrounding region in the 3xCTS-APEX2 line in order to identify the right labelling conditions. n= 1515 ± 347 cilia quantified per condition, each point represents one cilium. **** $p < 0.0001$ by Kruskal–Wallis test. **(C)** Example of Scarlet+ cilia and **(D)** surrounding region of each cilium segmentation using the Harmony software. **(E)** Immunofluorescence images of labelled and mock 3xCTS-APE2 hypothalamic neurons stained for neuronal (MAP2) and hypothalamic (POMC) markers. Scale bars represent 10 µm. Insets show channel-splitted cilia. **(F)** Quantification of DAPI+ cells expressing the 3xCTS construct. Number of Scarlet+ objects were counted and normalised to the number of nuclei. Each dot represents a single field of view (n =18) of 1 technical replicate. Bars indicate mean + SEM. **(G)** Western blot analysis of IFT88 and IFT57 in proteins extracted from 3xCTS-APEX2, TM-APEX2 or WT hypothalamic neurons after APEX2-mediated labelling (+) or mock labelling (-). **(H-I)** Quantifications of the signal intensity of IFT57 (G) and IFT88 (H) normalised to total protein stain in the eluate samples. ** $p < 0.0021$, *** $p < 0.0002$ and **** $p < 0.0001$ by Kruskal–Wallis test.

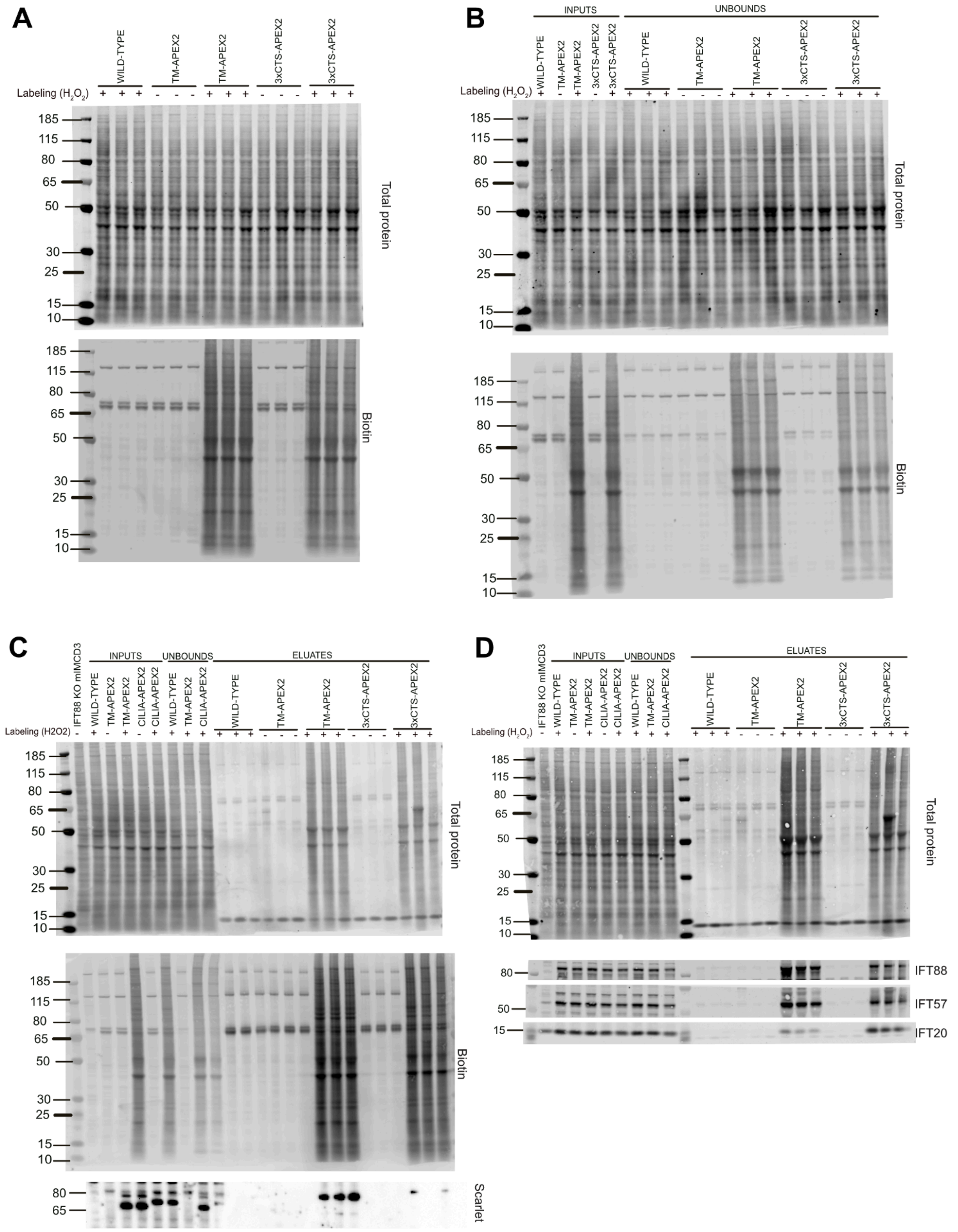


**Figure S2.** Original uncropped western blots of proteins extracted from 3xCTS-APEX2, TM-APEX2 or WT hypothalamic neurons after APEX2-mediated labelling (+) or mock labelling (-), run in triplicates. **(A)** Proteins were collected immediately after cell lysis (inputs). **(B)** Unbound material, that represents everything that did not bind to the chromatography column, was compared to inputs. **(C)** Biotinylated proteins after purification by streptavidin chromatography (eluates) were detected by streptavidin-HRP (Biotin) and compared to inputs and unbounds. **(D)** Other proteins after purification were detected by specific antibodies and compared to inputs and unbounds. Protein sizes are in kDalton.

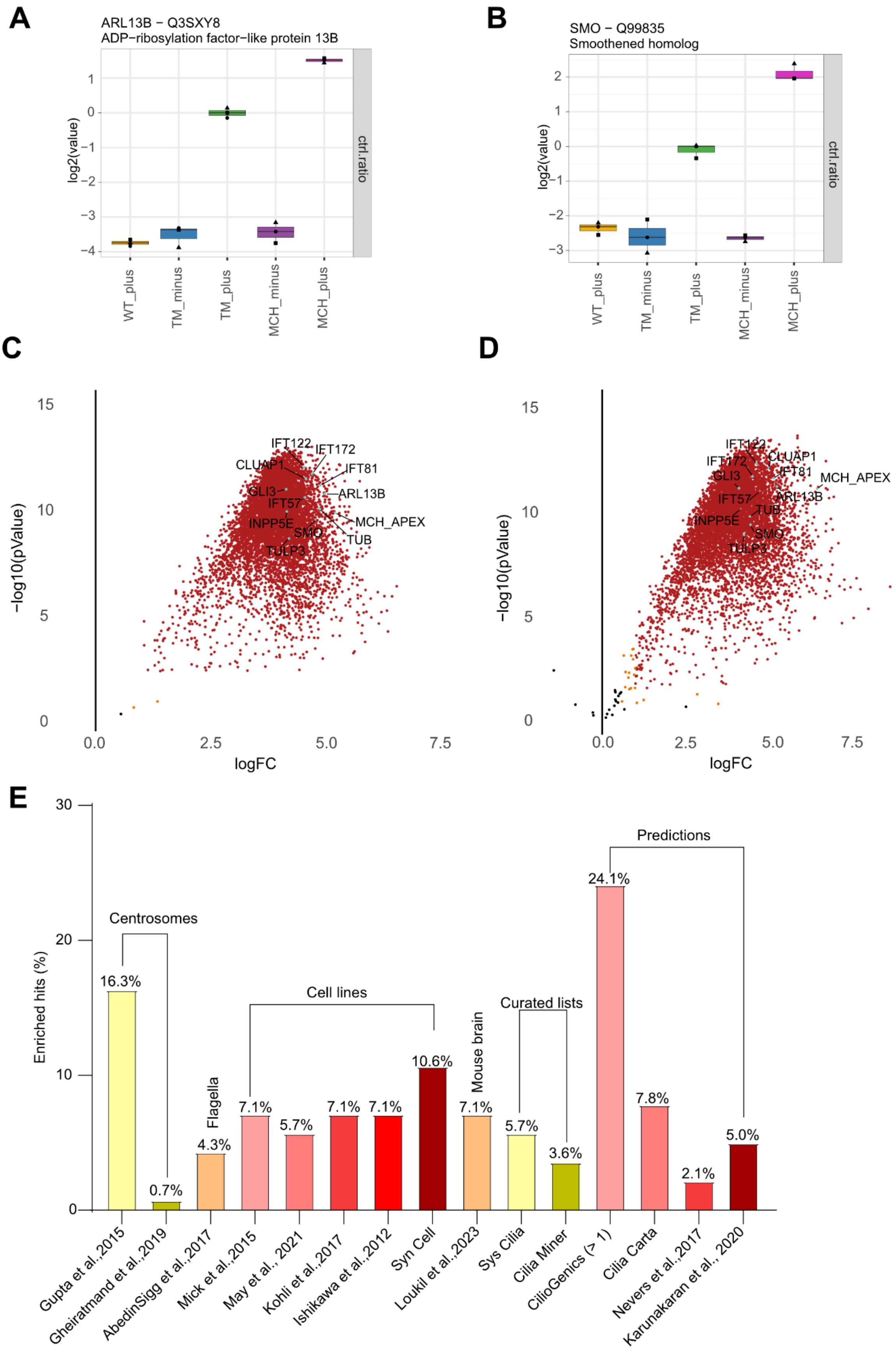
A
ARL13B − Q3SXY8
ADP−ribosylation factor−like protein 13B
log2(value)
ctrl.ratio
WT_plus
TM_minus
TM_plus
MCH_minus
MCH_plus
B
SMO − Q99835
Smoothened homolog
log2(value)
ctrl.ratio
WT_plus
TM_minus
TM_plus
MCH_minus
MCH_plus
C
−log10(pValue)
logFC
IFT122
IFT172
CLUAP1
IFT81
GLI3
ARL13B
IFT57
MCH_APEX
INPP5E
SMO
TUB
TULP3
D
−log10(pValue)
logFC
IFT122
IFT172
CLUAP1
IFT81
MCH_APEX
GLI3
ARL13B
IFT57
TUB
INPP5E
SMO
TULP3
E
Enriched hits (%)
Centrosomes
Flagella
Cell lines
Mouse brain
Curated lists
Predictions
16.3%
0.7%
4.3%
7.1%
5.7%
7.1%
7.1%
10.6%
7.1%
5.7%
3.6%
24.1%
7.8%
2.1%
5.0%
Gupta et al.,2015
Gheiratmand et al.,2019
AbedinSigg et al.,2017
Mick et al.,2015
May et al., 2021
Kohli et al.,2017
Ishikawa et al.,2012
Syn Cell
Loukil et al.,2023
Sys Cilia
Cilia Miner
CilioGenics (> 1)
Cilia Carta
Nevers et al.,2017
Karunakaran et al., 2020

**Figure S3. (A,B)** ARL13B and SMO protein abundance in WT, 3xCTS (MCH) and TM control samples labelled (plus) or mock (minus) with APEX2. N =3 and log2 Value intensities of each replicate are normalised to control (TM plus). **(C,D)** Volcano plots of statistical significance (pValue) versus protein enrichment (FC) in 3xCTS-APEX2 labelled (plus) compared to mock labelled (minus) samples (A) and compared to WT labelled (plus) samples (B). **(E)** The list of enriched hits was compared to lists of proteins published in different studies. % represents overlaps of the datasets with the enriched hits. Gupta et al. 2015, Gheiratmand et al. 2019 and Syn Cell (Y.-D. Chen et al. 2023) datasets were derived from human cell lines; Sigg et al. 2017 from sea urchins, sea anemones and choanoflagellates; Mick et al. 2015, May et al. 2021, Kohli et al. 2017, and Ishikawa et al. 2012 from mouse cell lines; Loukil et al. 2023 from mouse brains. Sys cilia (Vasquez, van Dam, and Wheway 2021) and Cilia Carta (van Dam et al. 2019) are curated lists from different species, while CilioGenics (Pir et al. 2024) includes only human genes. CiliaMiner (Turan et al. 2023) is a database of human ciliopathies genes, Nevers et al. 2017 includes predicted orthologs of human protein-coding genes and Karunakaran et al. 2020 represents bioinformatic prediction of interaction partners of ciliary genes.

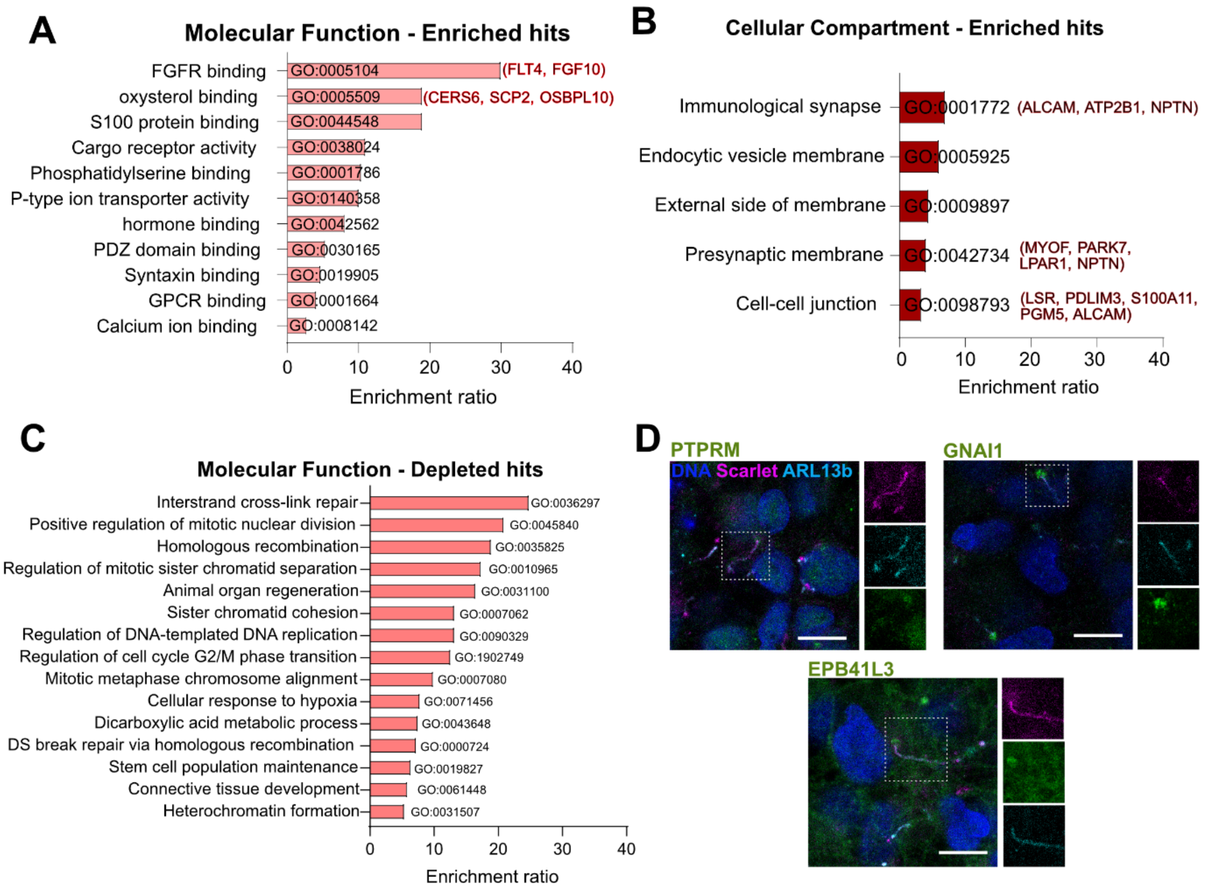


**Figure S4. (A-B)** GO term enrichment analysis for molecular function (A) and cellular compartment (B). Between brackets some example proteins of the term. **(C)** GO term enrichment analysis for molecular function of proteins enriched in TM control and depleted in cilia. **(D)** Immunofluorescence stainings for some of the candidates showed localisation to the cilium base. PTPRM: Receptor-type tyrosine-protein phosphatase mu; GNAI1: G Protein Subunit Alpha I1; EPB41L3: Erythrocyte Membrane *Protein* Band 4.1 Like 3. Scale bars represent 10 µm. Insets show channell-splitted cilia.

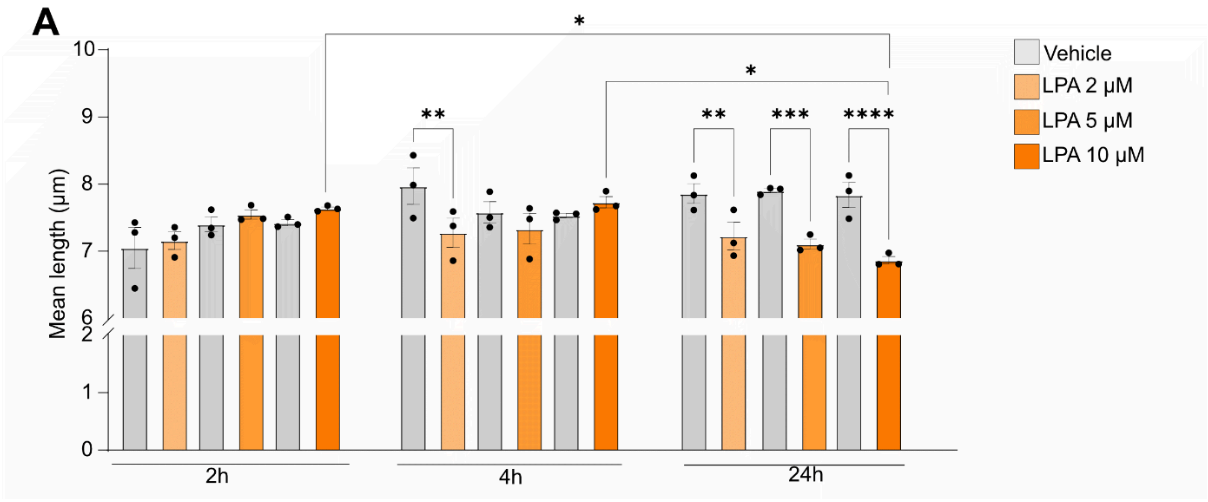


**Figure S5. (A)** Dose-response to LPA at different timepoints (2, 4, 24h) in hiPSC-derived hypothalamic neurons expressing the 3xCTS-APEX2 construct. Mean length is shown per replicate, N = 3. * $p < 0.0332$, ** $p < 0.0021$, *** $p < 0.0002$, **** $p < 0.0001$ by Kruskal–Wallis test.
